# Chimpanzees communicate to coordinate a cultural practice

**DOI:** 10.1101/2021.03.22.436386

**Authors:** Zoë Goldsborough, Anne Marijke Schel, Edwin J. C. van Leeuwen

## Abstract

Human culture thrives by virtue of communication, yet whether communication plays an influential role in the cultural lives of other animals remains understudied. Here, we investigated whether chimpanzees use communication to engage in a cultural practice by analyzing grooming handclasp (GHC) interactions – a socio-cultural behavior requiring inter-individual coordination for its successful execution. Previous accounts attributed GHC initiations to behavioral shaping whereby the initiator physically molds the partner’s arm into the desired GHC posture. Using frame-by-frame analysis and matched-control methodology, we find that chimpanzees do not only shape their partner’s posture (22%), but also use gestural communication to initiate GHC (44%), which requires an active and synchronized response from the partner. Moreover, in a third (34%) of the GHC initiations, the requisite coordination was achieved by seemingly effortless (i.e., no shaping or communication) synchrony. Lastly, using a longitudinal approach, we find that communication occurs more frequently than shaping in experienced dyads and less in mother-offspring dyads. These findings are consistent with the theory of ontogenetic ritualization, thereby reflecting the first documentation of learned communication in a cultural context. We conclude that chimpanzees show situation-contingent interactional flexibility in the socio-cultural domain, opening the possibility that the interplay between communication and culture is rooted in our deep evolutionary history.

## Introduction

Human culture is catalyzed by many forms of communication, ranging from active teaching to subtle cue-responding during turn-taking (Enfield and Levinson, 2006; Tomasello, 2010). By means of communication, information – including cultural knowledge – can spread efficiently, resulting in a bolstering of the within-group homogeneity and between-group heterogeneity typical of cultural variation (Boyd and Richerson, 1985; Gergely and Csibra, 2005). While pivotal to human culture, it is currently unknown whether non-human animal culture (henceforth ‘animal culture’) is similarly guided by communication. Animal culture has been defined in many ways (McGrew, 1998; Whiten and van Schaik, 2007), yet a shared feature across all definitions is that the corresponding behavior needs to be transmitted via social learning, which is the process by which individuals obtain information through observation or interaction with others or their products (Heyes, 1994). Moreover, many scholars adhere to the definitional criterion that such social learning processes ought to lead to group-specific behaviors (Hoppitt and Laland, 2013). Following this definition, there is currently little doubt that many animal species possess, and live by, cultural traditions (Allen, 2019; Aplin, 2019; Whiten, 2019).

However, an outstanding question related to animal culture concerns the specific transmission mechanisms of their traditions. How exactly does acquired behavior spread through groups of animals such that it becomes culture? An impressive body of experimental work has shown that animals are socially facilitated into new behavioral variations (Reader and Biro, 2010) and may even copy others by means of emulation/imitation – the processes by which observers learn from the consequences of others’ behavior, or from their very bodily movements, respectively [(Hoehl et al., 2019; Horner and Whiten, 2005; Whiten et al., 2009), cf. (Tennie et al., 2020)]. Additionally, there is a large body of evidence from the wild indicating that animals establish cultural traditions, ranging from vocal dialects in birds and whales (Aplin, 2019; Noad et al., 2000; Zandberg et al., 2021) to arbitrary conventions in meerkats and primates (Huffman et al., 2008; Perry et al., 2003; Thornton et al., 2010).

Yet, what we currently do not know is how animals transmit cultures that solely exist by means of the interactions between individuals (henceforth ‘cultural interactions’). For instance, in humans, there are many cultural behaviors that ‘take two to tango’, such as the tango itself, but also many dyadic encounters like greeting exchanges and conversing (Enfield and Levinson, 2006; Stivers et al., 2009). Do animals have similar cultural interactions? And if so, how are they instigated and maintained? Moreover, we do not yet know whether animals actively communicate to transmit these interactive cultures. To our knowledge, most of the documented examples of animal culture concern instances in which an observer learns from an otherwise passive other. In other words, individual A copies the behavior of individual B without individual B actively transmitting its cultural knowledge. The more active forms of cultural transmission are embodied in human pedagogy and teaching (Gergely and Csibra, 2005), yet while there are several indications that animals may teach as well (Boesch, 1991; Musgrave et al., 2016; Thornton and Raihani, 2010), the consensus lies, at least in chimpanzees, with a minimal account of teaching in which an active role of the purported teacher has yet to be identified (Lonsdorf, 2006; Moore, 2013).

Here, we investigate an enigmatic cultural interaction in chimpanzees – the grooming handclasp (McGrew and Tutin, 1978) – to test whether chimpanzees may actively communicate to coordinate these interactions. In a grooming handclasp (henceforth ‘GHC’) two partners extend one of their arms overhead and clasp each other’s extended hand at the palm, wrist, or forearm, while grooming each other with the other arm (McGrew and Tutin, 1978; Nakamura and Uehara, 2004; van Leeuwen et al., 2012; Figure 1). While the cultural nature of GHC has been firmly established by reports on inter-group differences in the form and frequencies of practices (McGrew et al., 2001; Nakamura, 2002; van Leeuwen et al., 2017, 2012; Wrangham et al., 2016), little is known about the ways in which chimpanzees coordinate the execution of this cultural interaction, other than one individual (i.e., the initiator) physically shaping the body of the envisioned partner into the GHC posture (de Waal and Seres, 1997). To learn more about this coordination process, we studied the behaviors associated with the onset of GHCs by means of frame-by-frame analysis in a group of semi-captive chimpanzees in Zambia and compared the observed behaviors with matched-control windows (i.e., social grooming events of the same partners *without* GHC (*sensu* de Waal and Yoshihara 1983)). Consistent with reports evidencing intentional communication in social contexts like joint travel and play (Fröhlich et al., 2016a, 2016b), we hypothesized that chimpanzees would not merely shape their partner into the typical handclasp posture (de Waal and Seres, 1997), but actively communicate to coordinate their cultural practice. Finding support for this hypothesis would provide the first evidence for non-human animals to organize their cultural lives by means of proactive communication, akin to the human species.

**Figure 1.**
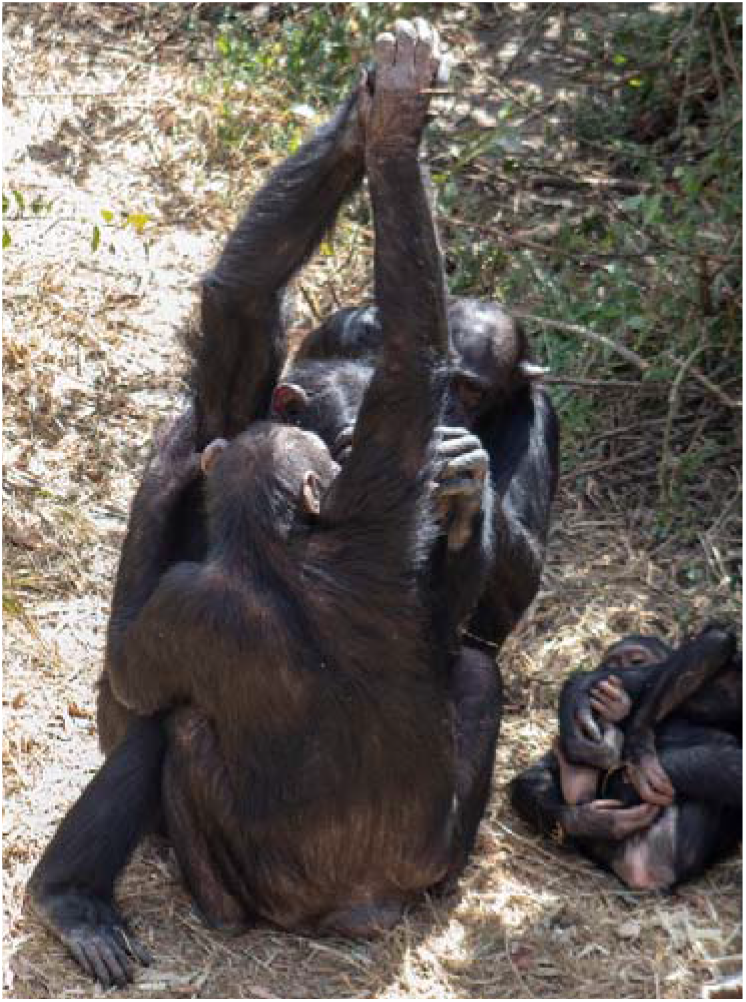
Debbie (back) and Tess (front) engaged in a palm-to-palm grooming handclasp (back-view: photo by Zoë Goldsborough).

## Methods

### Subjects

Subjects were 52 semi-wild chimpanzees (Table S1) at the Chimfunshi Wildlife Orphanage, Zambia, a sanctuary where chimpanzees live in Miombo woodland enclosures (size = 65 ha) where they can nest and forage independently but do receive daily feedings (van Leeuwen et al., 2012). GHC has been frequently observed in the group for over 12 years (van Leeuwen, 2021). The study was approved by the Chimfunshi Research Advisory Board (permit: CWOT_ 2019C039) and conformed to the nationwide legal requirements. Chimfunshi is accredited by PASA and adheres to the rules and regulations with respect to animal care and management as stipulated by the Zambia Wildlife Authority.

### Collection and Coding

Data were collected by ZG from 23-03-2019 to 04-06-2019 between 8am-4pm with handheld digital video cameras (Panasonic HDC-HS100). To capture GHC initiations, filming commenced as soon as two individuals approached one another. Filming continued if the individuals started social grooming (uni- or bi-directional) and lasted until they (a) had stopped grooming for over 30 seconds, (b) started grooming another individual, or (c) physically separated. A grooming bout was defined as running from the start of grooming until the moment one of the aforementioned ending conditions were met. A bout was considered a GHC-bout if it contained one or multiple GHCs, and a regular grooming bout if no GHCs occurred.

GHC-bouts had either a *side* (optimal) or *back* (sub-optimal) view. If a 10sec pre-handclasp (PH) social grooming window was available before the first GHC in the bout (see Figure 2), we used the video for analysis. We only analyzed the initiation of the first GHC in GHC-bouts, because previous GHCs could possibly function as signals for subsequent GHCs. The start of a GHC was defined as the instance of handclasp above face level; the end was defined as the instance that physical contact of the arms was broken. A Matched-Control (MC) period (see Figure 2) was analyzed to enable comparison of individual initiation behaviors across conditions (de Waal and Yoshihara, 1983). The MC-period was defined as a 10sec-window minimally 10sec after the last GHC occurrence in the bout, in which the same individuals had to be positioned in the same relative positions as during the GHC, while still engaging in social grooming. Additionally, initiations of regular, non-GHC-bouts were opportunistically recorded (*n_side + back_*=23) to identify behaviors used in the initiation of regular social grooming bouts.

**Figure 2.**
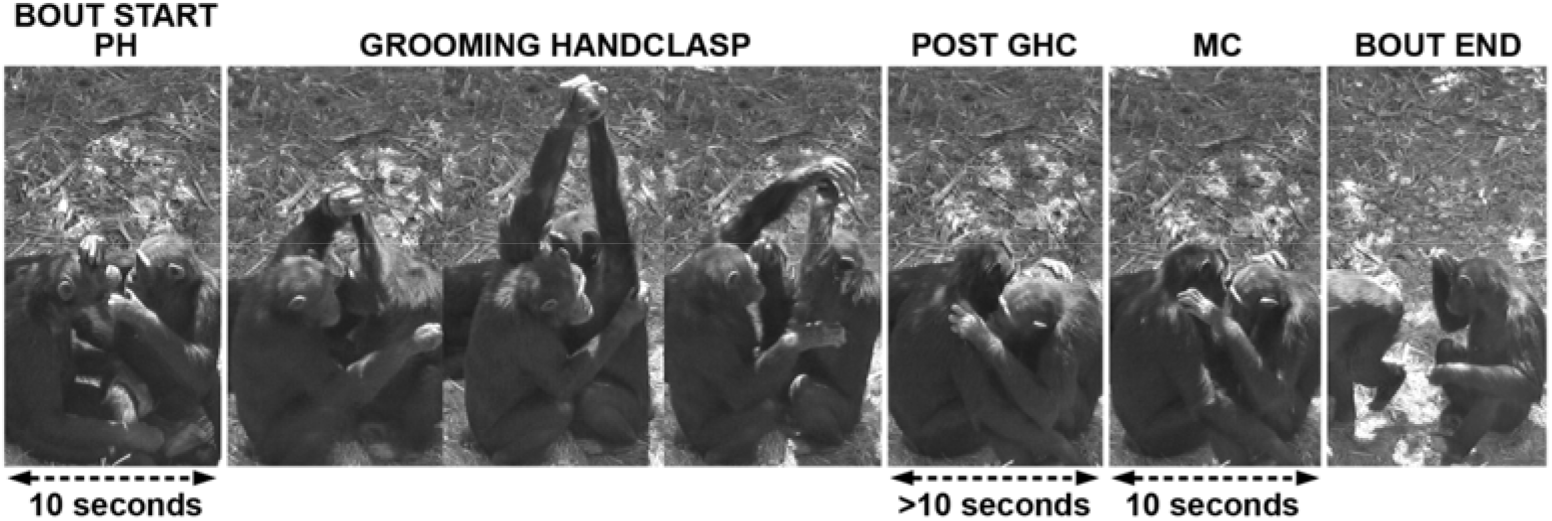
Side-view of GHC bout including the identified pre-handclasp (PH) and matched-control (MC) period. The PHs and MCs were chosen to exactly match in terms of individuals, bodily positioning and activities (grooming) in order to identify the mechanisms by which GHC is initiated.

Videos were scored in ELAN (Wittenburg et al., 2006) and behaviors were coded based on preliminary screening of the videos and established chimpanzee ethograms (Nishida et al., 1999) (see Table S2 and SI videos). A subset of 20% of the data was coded by two further observers to establish inter-rater-reliability (IRR). Mean dyadic agreement was 0.833 for coding behaviors (range 0.81-0.89), and 0.973 for identifying the initiating individual (range 0.89-1; see SI for details).

### Analyses

To determine the mechanisms underlying GHC initiation, we investigated the occurrences of 10 selected chimpanzee behaviors (Table S2 and Table S3) in a comparison between pre-handclasp (PH) periods and their matched-control (MC) windows in video-recorded GHC bouts with the optimal vantage point (*side*-view, *n*_ph-mc_=94; *n*_ind_=33, n_dyads_=48, see Figure 2). In this sample, the mean number of bouts that an individual was involved in was 5.7 (range 1-28), with 8 individuals only involved in one bout. All analyses were done in R v. 4.0.3 (R Core Team, 2020). When necessary, non-parametric statistics were applied, including Bonferroni-Holm corrections for multiple testing (Holm, 1979).

In the PH-MC comparison, we only analyzed those behaviors that occurred ≥5 times in the PH and MC of GHC-bouts (see Table S3). Furthermore, two behaviors (“nosewipe” and “self-scratch”) were excluded from this analysis, as they are known self-directed behaviors linked to increased arousal (Aureli and de Waal, 1997; Leavens et al., 2004) and, as such, may also be produced in other contexts besides initiating a GHC interaction (though they may be inadvertent signals, revisited below). The behavior “torso” was also excluded as it is potentially an artifact of chimpanzees turning towards their partner as a necessary prelude for a grooming interaction. Additionally, given that social grooming occurred in both PH and MC windows by definition (see Figure 2 and “Collection and Coding”), we did not consider the grooming behaviors themselves (see Table S3) as possible signals for GHC initiation. Based on our findings (see Results) and previous literature (e.g. de Waal and Seres, 1997), we classified the behaviors identified in the PH-MC comparison into three types of GHC initiations: *i*) Shaping, *ii*) Communication, and *iii*) Synchrony. We also conducted auxiliary analyses including *back*-view observations of GHC-bouts (*n*_total_=133, *n*_individuals_ =34, *n*_dyads_=57). In this larger sample, the mean number of bouts per individual was 7.7 (range 1-35), with 7 individuals only involved in one bout.

To assess the flexibility of and presence of elaboration in GHC initiations, we used all *side*-view GHC bouts regardless of the presence of MCs (*n*=114, *n*_individuals_ =34, *n*_dyads_=58). In this sample, the mean number of bouts an individual was involved in was 6.5 (range 1-24), with 8 individuals only involved in one bout. We considered the flexibility of GHC initiations by exploring variation in the start behavior of the initiator as well as variation in their behavioral sequences in the PH period. An individual was considered the initiator of a GHC bout when they were either *a*) the first one to produce a GHC-specific initiation behavior or *b*) in the absence of these behaviors, the first to raise their arm for the GHC. Note that 7 bouts have been dropped from this sample, as these were initiations where the initiator did not show any behaviors before raising their arm. We define elaboration as the use of new or additional behavior after an initial behavior (Leavens et al., 2005).

Lastly, we examined what factors could account for how a dyad initiated a GHC. Mothers play a more active role in the acquisition of the GHC by their offspring than other group members (van Leeuwen et al. 2012; Nakamura, 2002), largely through physically shaping their offspring in the GHC posture. Thus, we expect that GHCs between mother-offspring dyads are more likely to be initiated via physical shaping than GHCs between dyads that are not mother-offspring. Additionally, dyads with more experience engaging in GHC with each other could be better at coordinating the GHC interaction as a result of many repeated interactions. As such, experienced dyads might be less likely to initiate GHCs via physical shaping than inexperienced dyads. To test this, we employed a Bayesian categorical (multinomial) model on all *side*-view PH-MC bouts (*n*= 94). The outcome variable was the type of GHC initiation observed in a bout (Shaping, Communication, or Synchrony, with Shaping as reference category), and the predictor variables were *i)* days of experience (days since this dyad’s first GHC, based on a longitudinal dataset of opportunistic GHC observations between 2007-2019, standardized such that 0 is no experience and 1 the highest number of days in the sample) and *ii)* whether the dyad was a mother-offspring dyad or not. To account for dyadic or initiator preferences for specific initiation types, we included dyad and initiator ID as random effects. This model was fitted with the brms package v. 2.16.1 (Bürkner, 2017). We set mild regularizing priors – normal(0,1) for estimates and exponential(1) for standard deviations – and performed a prior predictive simulation to visualize the priors. For running the final model, we used four chains, 10,000 iterations, and a credibility interval of 0.95. Our model was stable with large effective sample sizes (Bulk_ESS and Tail_ESS over 1000) and Rhat values equal to 1.

## Data availability statement

Code and data used for analyses are available via https://doi.org/10.5281/zenodo.6805926.

## Results

We observed a total of 548 GHC initiations (of which 133 were recorded on video) during the 5-month study period (Table S1). Of the 52 chimpanzees in the group, 39 individuals (including all adults) were observed to engage in GHC at least once.

### GHC-specific behaviors

Seven behaviors were observed more frequently in the PH compared to the MC context (Wilcoxon signed-rank: all *p*<0.04, Holm-corrected; see Table S4). These behaviors were thus considered to be potential mechanisms leading to GHC interactions. Due to their physical nature, two of these behaviors (“elbow hold” and “hand grab”) corresponded to the documented practice of *shaping* (de Waal and Seres 1997; e.g., Video S11), while the remaining behaviors (“elbow touch”, “hand touch”, “head move”, “head touch”, and “hold”) lacked any prolonged physical contact with the partner and were thus considered to be potential *communicative gestures* (Liebal and Call 2012; e.g., Video S12). Auxiliary analyses including the sub-optimal *back*-view bouts supported our main analyses (all 7 initiation behaviors significantly more present in PH than MC, all *p*<0.02, Holm corrected; see Tables S3 and S4).

Gestures are here defined as bodily actions directed at a conspecific that are mechanically ineffective and potentially result in a voluntary response from the recipient (Pika, 2008; Liebal and Call 2012). The five potentially communicative GHC-initiation behaviors complied with this definition in being mechanically ineffective bodily actions resulting in voluntary GHC responses. Only for “head move” we could envision partly co-variation with other behaviors such as changing grooming posture or redirecting attention. “Elbow touch”, “hand touch”, and “head touch” involved targeted physical contact from actor to recipient and were thus directed at a conspecific, and during “hold” and “head move” signalers faced their recipient in 100% of observed instances (*n*=31 and *n*=18, respectively). 14 of 15 individuals performing more than one GHC initiation showed variation in the start behavior (i.e., the 7 GHC-specific behaviors determined above) of their initiation sequences (Binomial test: *p*<0.001; also see Figure 3 and associated R-code).

**Figure 3.**
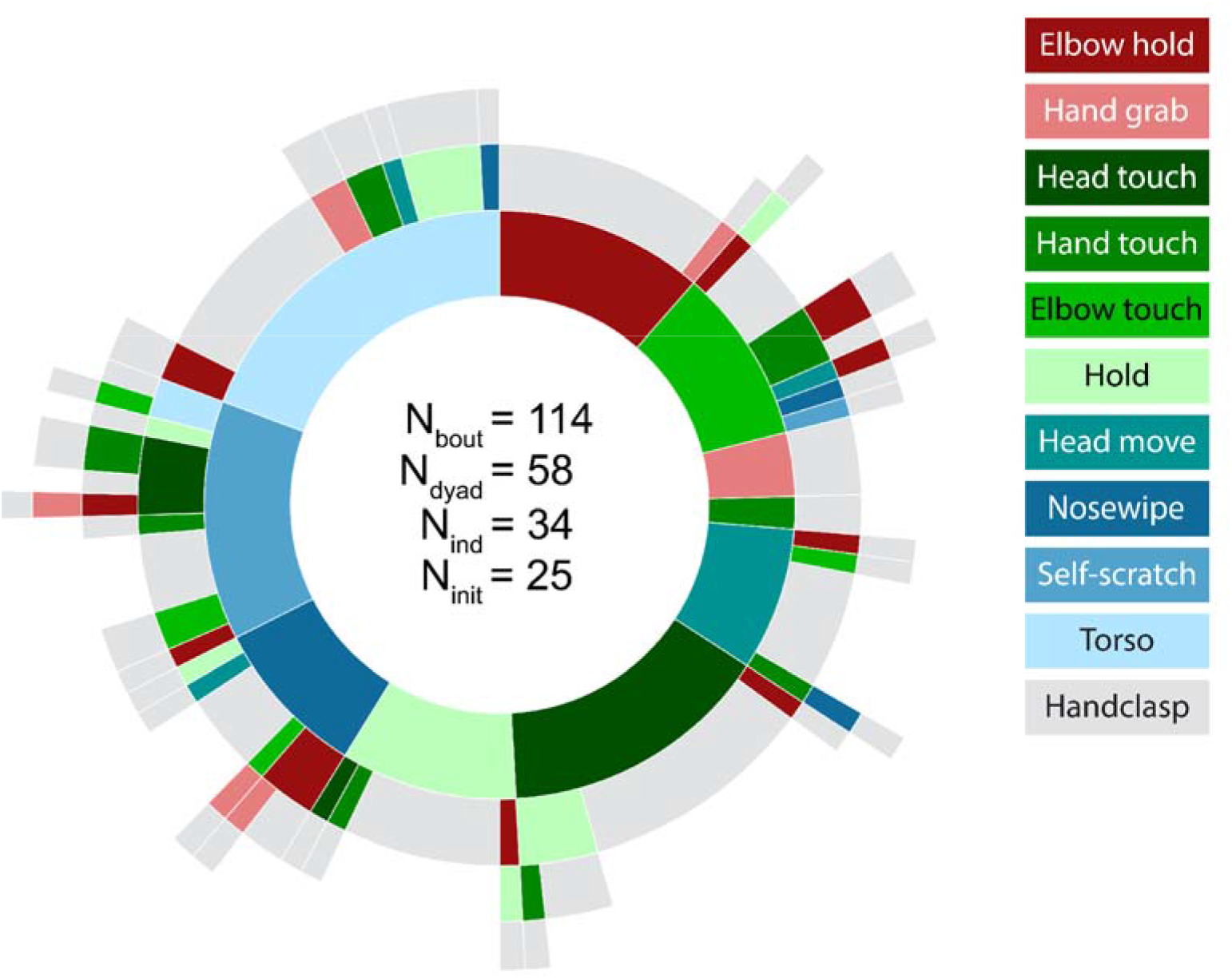
Sunburst (Bostock et al., 2020) showing behavioral sequences (*n*=114) by the initiator (*n* = 25) leading to GHC. Starting behaviors are depicted in the inner colored circle, with the grey outer circle being the endpoint of the sequence (i.e., GHC). In order to consider the full flexibility of all types of GHC initiations, we also included the three “synchrony” behaviors (“nosewipe”, “self-scratch”, and “torso”). * Interactive version available as Figure S13.

Moreover, the gestures were produced in a goal-directed way, as indicated by the occurrence of *elaboration* in 29% of the cases where an initial gesture failed to initiate a GHC (*n*=20 out of the 69 instances where gestures were used in the initiation, see Figure 3 and details below). Elaboration occurred after an average response waiting time of ~0.5s and took the form of another gesture (*n*=11), a shaping behavior (*n*=4), or a combination of both another gesture and a shaping behavior (*n*=5) before the GHC finally commenced. Taken together, these observations show that chimpanzees are capable and determined to (re-)transmit their motivation to engage in GHC when needed.

### GHC initiation types

In general, of the 94 PH/MC comparison bouts, 21 (22%) contained either one or both shaping behaviors (“elbow hold”, “hand grab”), 41 (44%) contained one or more of the five communicative gestures (“elbow touch”, “hand touch”, “head move”, “head touch”, “hold”) and no shaping behaviors, and 32 (34%) contained neither shaping behaviors nor potentially communicative behaviors. We labeled the third type of GHC initiation as “synchrony”, as the individuals appeared to commit to the GHC near-simultaneously. If any behavior was scored during the PH window in the synchronous GHCs these were either the previously mentioned self-directed behaviors, namely “self-scratch” and “nosewipe”, or “torso”. These behaviors might signal to a partner that an individual is in a high state of arousal (Leavens et al., 2004) or on the verge of initiating a grooming interaction (Wilke et al., 2022), which the partner could respond to by raising their arm for a GHC. The reason that we do not consider these behaviors as GHC-specific signals, however, is that these three behaviors were relatively frequent in the initiation of regular grooming bouts as well, whereas most of the communicative signals were entirely absent in the regular grooming bouts (see Table S3).

We found that the probability of a dyad initiating a GHC bout via communication rather than shaping increases when the dyad has more GHC experience (Table 1 and Figure 4a). Here, 91% of the posterior distribution lies above zero, which indicates that a positive relationship is expected to be observed 91% of the time. Additionally, mother-offspring dyads appear less likely to initiate a bout via communication (compared to shaping) than non- mother-offspring dyads (Table 1 and Figure 4b), with a probability of observing a negative relationship 89% of the time.

**Table 1.**
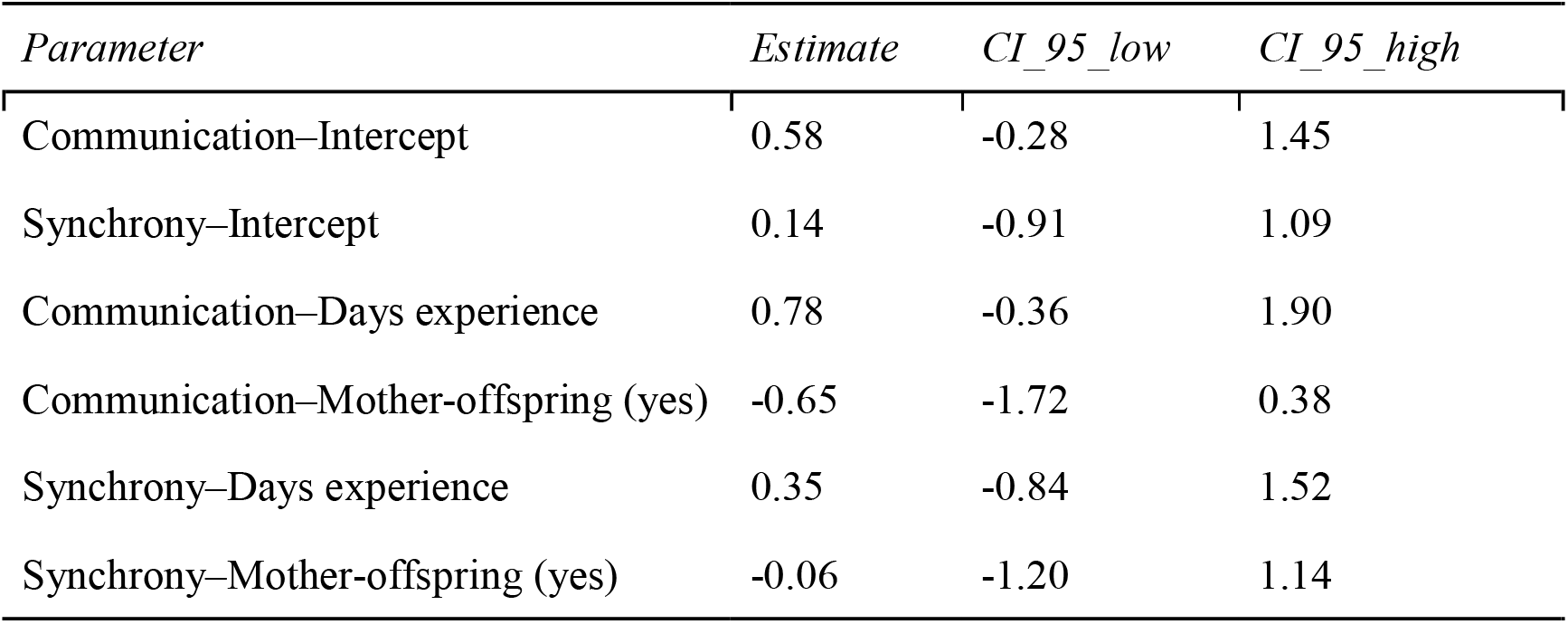
Results of multinomial model testing for the effect of GHC experience and mother-offspring kinship on type of GHC initiation. The parameter estimates are on the logit scale and are in relation to the reference category of the model, which is Shaping.

| *Parameter* | *Estimate* | *CI_95_low* | *CI_95_high* |
| --- | --- | --- | --- |
| Communication–Intercept | 0.58 | -0.28 | 1.45 |
| Synchrony–Intercept | 0.14 | -0.91 | 1.09 |
| Communication–Days experience | 0.78 | -0.36 | 1.90 |
| Communication–Mother-offspring (yes) | -0.65 | -1.72 | 0.38 |
| Synchrony–Days experience | 0.35 | -0.84 | 1.52 |
| Synchrony–Mother-offspring (yes) | -0.06 | -1.20 | 1.14 |

**Figure 4.**
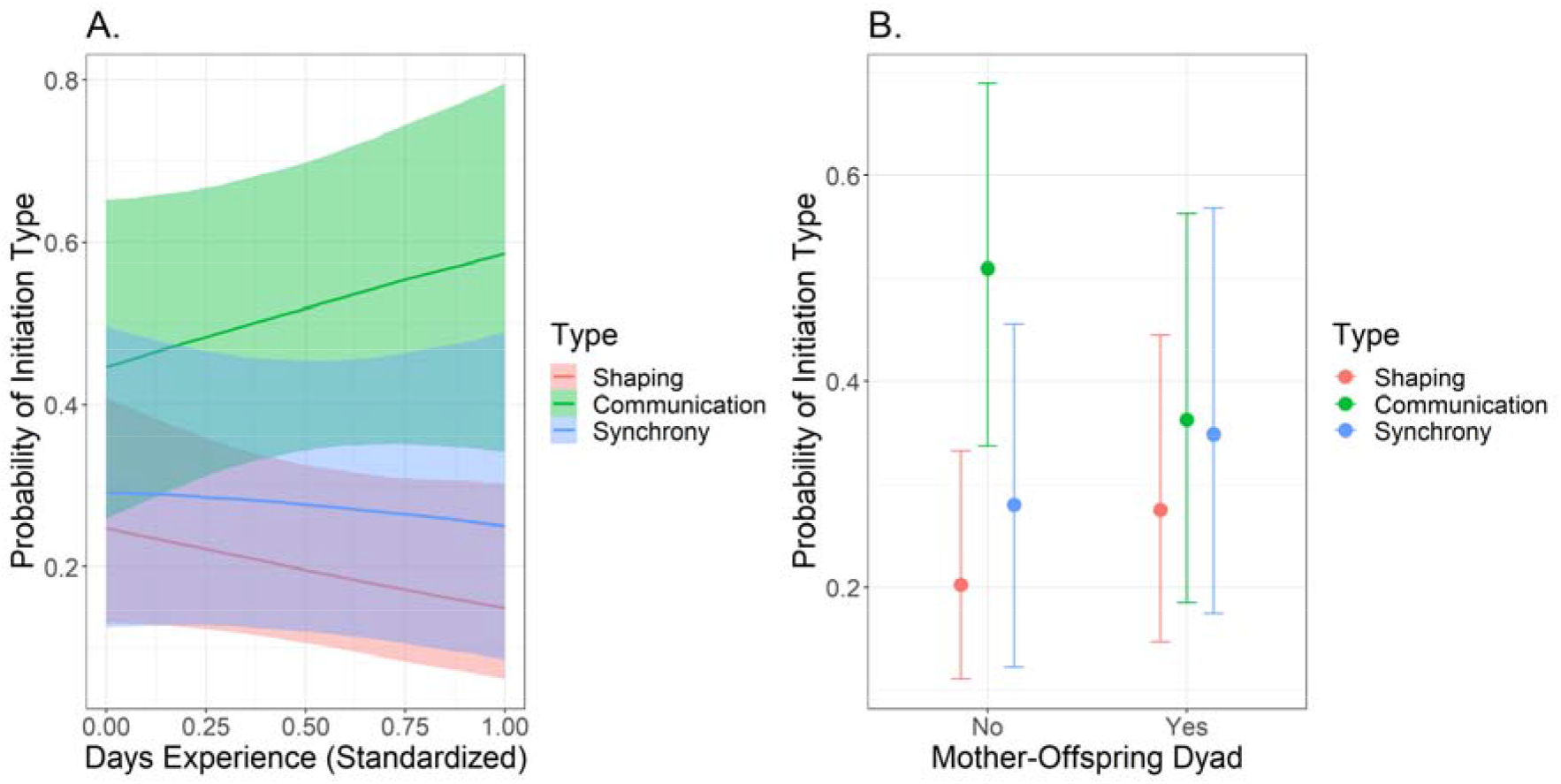
Probability of GHC initiation type in relation to a) dyadic days of experience with GHC and b) whether the dyad is a mother-offspring dyad or not. The lines in a) and dots in b) indicate the model fit of the multinomial model in Table 1, with the shaded areas corresponding to the 95% credibility interval.

## Discussion

In this study we set out to test whether communication may play a role in facilitating cultural practices in animals other than humans. Human culture is vastly nurtured by means of communication (e.g., verbal instructions on how to behave in the classroom), but whether such influences exist in the animal kingdom remains to a large part unknown. Here, we show that chimpanzees communicate to engage in one of their most enigmatic socio-cultural practices, the grooming handclasp (McGrew and Tutin, 1978). In addition to physically shaping a partner into the GHC position – the hitherto only explanation for the coordination of the GHC – we identify goal-directed gestural communication as an additional mechanism by which chimpanzees initiate and coordinate their culturally bestowed handclasps.

While it is known that chimpanzees intentionally communicate to entice group members into desired responses (Hobaiter and Byrne, 2011; Fröhlich et al., 2016a, 2016b), to date, such communication has not been reported in the context of cultural practices. The grooming handclasp is arguably one of the most convincing examples of animal culture for at least three reasons: (1) there is substantial variation with respect to its prevalence in groups of both chimpanzees and bonobos (McGrew and Tutin, 1978; Whiten et al., 1999), (2) within the chimpanzee groups that engage in the GHC, there is substantial and stable variation in the specific technique preferred (McGrew et al., 2001; Nakamura, 2002; van Leeuwen, 2021; van Leeuwen et al., 2012), and (3) the GHC does not require environmental input, making it a behavioral phenomenon that is less dependent on the local availability of materials than most other cultural traditions in chimpanzees (Kalan et al., 2020; Whiten et al., 1999). The latter reason makes it less probable that the GHC is in fact a behavior that is instigated and formed by non-cultural determinants (van Leeuwen, 2021), much like vocal cultures in birds (Aplin, 2019) and cetaceans (Garland and McGregor, 2020). Unlike vocal cultures in birds and cetaceans, however, in the chimpanzees’ handclasp case, the (non-vocal) communication is not cultural in itself, but used to guide the enactment of their cultural practice. As such, irrespective of the chimpanzees’ being aware of the cultural nature of the GHC, they actively engage in a socio-cultural behavior by a variety of social means, including communication.

Previously, it was known that chimpanzees solve the coordination problem inherent to handclasping by means of physically shaping the desired partner into the typical A-frame posture of the GHC (de Waal and Seres, 1997). Here it is important to note that this mechanism requires one adamant individual and at most a passive, yet non-declining partner. The ensuing process is consistent with described patterns of transmission: for a long period of time, typically there is one eager individual in a group who initiates most, if not all GHC interactions (e.g., chimpanzee “Georgia” in (Bonnie and de Waal, 2006; de Waal and Seres, 1997)). However, this propagation mechanism (i.e., the proactively shaping of a willing partner) only covers the early stages of transmission – what mechanisms might sustain this socio-cultural tradition once there are more proficient group members? While shaping may still be at play, with two skilled and motivated partners, the GHC becomes more fluent and bidirectional. In other words, our finding that chimpanzees also use non-physical means to initiate a handclasp identifies an active partner response (otherwise the GHC would not ensue) and implicates a willingness from both partners to engage in this cultural practice.

In this light, it is worth highlighting that the active involvement of the partner complements the initiated GHC sequence by virtue of which the interaction may be considered as a joint action – at least in comparison to shaping interactions. With two voluntarily acting partners (i.e., one chimpanzee showing this by initiating, the other chimpanzee showing this by responding) in a tightly coordinated interaction (i.e., there is only a small window in spacetime where the arms can clasp), the GHC may offer a fruitful context in which to study joint commitment and perhaps even joint/shared intentionality. Future research in this domain may benefit by extending the scope of GHC scrutiny from initiations (this study) to interactions during and revolving around the ending of the interaction (Genty et al., 2020).

Our GHC analyses revealed a third coordination process which we coined “synchrony” due to its indiscernible execution. In these cases, the GHC coordination was fluent to the extent that no shaping or communication was required to accomplish it. We conjecture that perhaps repeated GHC engagement attunes partners’ behavior to the extent that a subtle indication, be it behaviorally (e.g., a nose wipe, a slight change in body orientation) or embodied by contextual factors (e.g., the order or duration of the ongoing grooming session), suffices to achieve coordination. Yet, our analyses did not indicate that more experienced dyads showed more “synchronous” GHC interactions. It may be that our sample was too dispersed to obtain a reliable indicator of “experience” (i.e., spanning relatively short observation windows between 2007-2019). Alternatively, it might be that our ethogram lacked the resolution to identify coordination behaviors that preclude the “synchronous” label. Finally, it may be that other factors than mere experience contribute to partners becoming fluent at achieving fine-tuned coordination, for instance individuals’ motivation and capacity to pick up on social cues.

The findings that *i*) communication is used more frequently as GHC initiation strategy than shaping in experienced dyads, and *ii*) shaping is more frequent in mother-offspring dyads than in non-mother-offspring dyads, are largely consistent with ontogenetic ritualization playing a role in the development of this communicative strategy (Tomasello and Call, 2007). This theory suggests that some behaviors can start to function as communicative signals through mutual anticipation of both partners following repeated shaping interactions. Inexperienced GHC dyads, such as a mother with younger offspring, may use physical shaping behaviors like “elbow hold” and “hand grab” to initiate a GHC. Over time, the offspring learns to anticipate the hold or grab and starts raising their arm at a touch of the elbow or hand, without requiring it to be held. However, not all gestures linked to the GHC initiation can be explained through ontogenetic ritualization: for instance, “hold” and “head touch” are unlikely to have arisen in this way, and many other gestures in the chimpanzee repertoire cannot be explained as such (Hobaiter and Byrne, 2011)◻.

If ontogenetic ritualization is the driving force of the development of social customs in chimpanzees, one would expect to find idiosyncratic gestures: different dyads employing different gestures for the same purpose. However, in the case of GHC, this would not be very probable since there are only limited ways to shape another individual’s body into the necessary A-frame posture typical of the GHC. Therefore, even following the theory of ontogenetic ritualization, within the GHC context, different dyads could end up using similar gestures to initiate a GHC, like “elbow touch” and “hand touch”. Of the remaining gestures, “hold” has previously been named as a behavior to invite another individual to handclasp in chimpanzees (Nishida et al., 2010) and bonobos (Fruth et al., 2006). The gesture “head touch” could be a soliciting act (Moore, 2013) to draw a partner’s attention and indicate an intent to groom or start an interaction. In the context of this specific group of chimpanzees, where GHC is so common that we rarely observe regular grooming bouts, “head touch” might have come to serve, fortuitously, as a signal for GHC initiation as this is the most likely grooming interaction.

Finally, our results provide new insights into how chimpanzees may coordinate their actions in general. The grooming handclasp is a social activity that requires coordination for successful execution. Chimpanzees cooperate (Mitani, 2009), but not much is known about the ways in which they coordinate their joint efforts (Duguid et al., 2020a). In experimental settings, some chimpanzees used location enhancing behaviors (e.g., bodily positioning, touching, peering) (Melis and Tomasello, 2019), or generic gestures (e.g., arm fling, clapping, banging on panels; Voinov et al. 2020) to recruit their conspecific partners for (re- engaging in) a cooperative task. While these behaviors can be interpreted as communicative acts to overcome coordination problems (for alternative interpretations, see Duguid et al., 2020a; Melis et al., 2018; Tennie et al., 2016), to date, it remains an interesting question why chimpanzees seem to use communication so little in contexts where it seems so obvious for humans (Duguid et al., 2020b).

Our findings show that chimpanzees *do* communicate to coordinate a naturally-occurring cultural practice. The socio-cultural interaction was not shaped by just one invested individual (Bonnie and de Waal, 2006; de Waal and Seres, 1997), but when the initiator communicated its desire to engage in the GHC (e.g., by holding out its flexed arm at face level in front of the desired partner), an active compliance in the form of a complementary action by the partner was required to accomplish the interaction. Thus, when communicated, a GHC initiation appeared to function as an invitation to join in a cultural practice. Following up on reports showing that chimpanzees and bonobos use communication during interactions like joint travel and social play (Fröhlich et al., 2016a; Heesen et al., 2017; Hobaiter and Byrne, 2014), we conclude that chimpanzees communicate to coordinate a cultural practice, and propose that our findings warrant future scrutiny of the grooming handclasp as a putative case of joint intentionality in great apes (Genty et al., 2020; Heesen et al., 2017, 2020).

## Supporting information

InteractiveSunburst

S1_Elbowhold

S2_Elbowtouch

S3_Handgrab

S4_Handtouch

S5_Headmove

S6_Headtouch

S7_Hold

S8_Nosewipe

S9_Selfscratch

S10_Torso

## Acknowledgments

We are grateful to the Chimfunshi Wildlife Orphanage Trust for their ongoing support of our research endeavors. In particular we wish to thank the staff of Chimfunshi, as well as Jake Brooker for his assistance during data collection and inter-rater-reliability.

## Compliance with Ethical Standards

The authors declare no conflict of interests to exist. The research on chimpanzees was strictly non-invasive/observational and adhered to the ethical guidelines provided by the Chimfunshi Research Advisory Board. All applicable international, national, and/or institutional guidelines for the care and use of animals were followed.

## Supplementary Information captions

**SI_1.** Supplementary information and tables, including subject demographics, the behavioral ethogram, frequency table of behavior, outcomes of the Wilcoxon-signed rank tests, posterior distributions of the multinomial model, and details on the inter-rater reliability calculation.

**Video S1**. Elbow hold.

**Video S2**. Elbow touch.

**Video S3**. Hand grab.

**Video S4**. Hand touch.

**Video S5**. Head move.

**Video S6**. Head touch.

**Video S7**. Hold.

**Video S8.** Nosewipe.

**Video S9.** Self-scratch.

**Video S10.** Torso.

**Video S11.** Video example of a grooming handclasp initiated with shaping behavior.

**Video S12.** Video example of a grooming handclasp initiated with gestural communication.

**Interactive Figure S13.** Interactive version of Figure 3.

## Supplementary Information

**Table S1.**
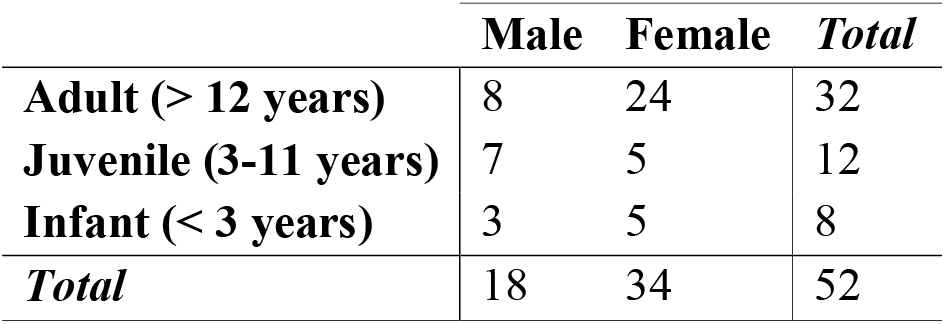
Subject demography Group 2 at Chimfunshi Wildlife Orphanage on 01-07-2019.

**Table S2.**
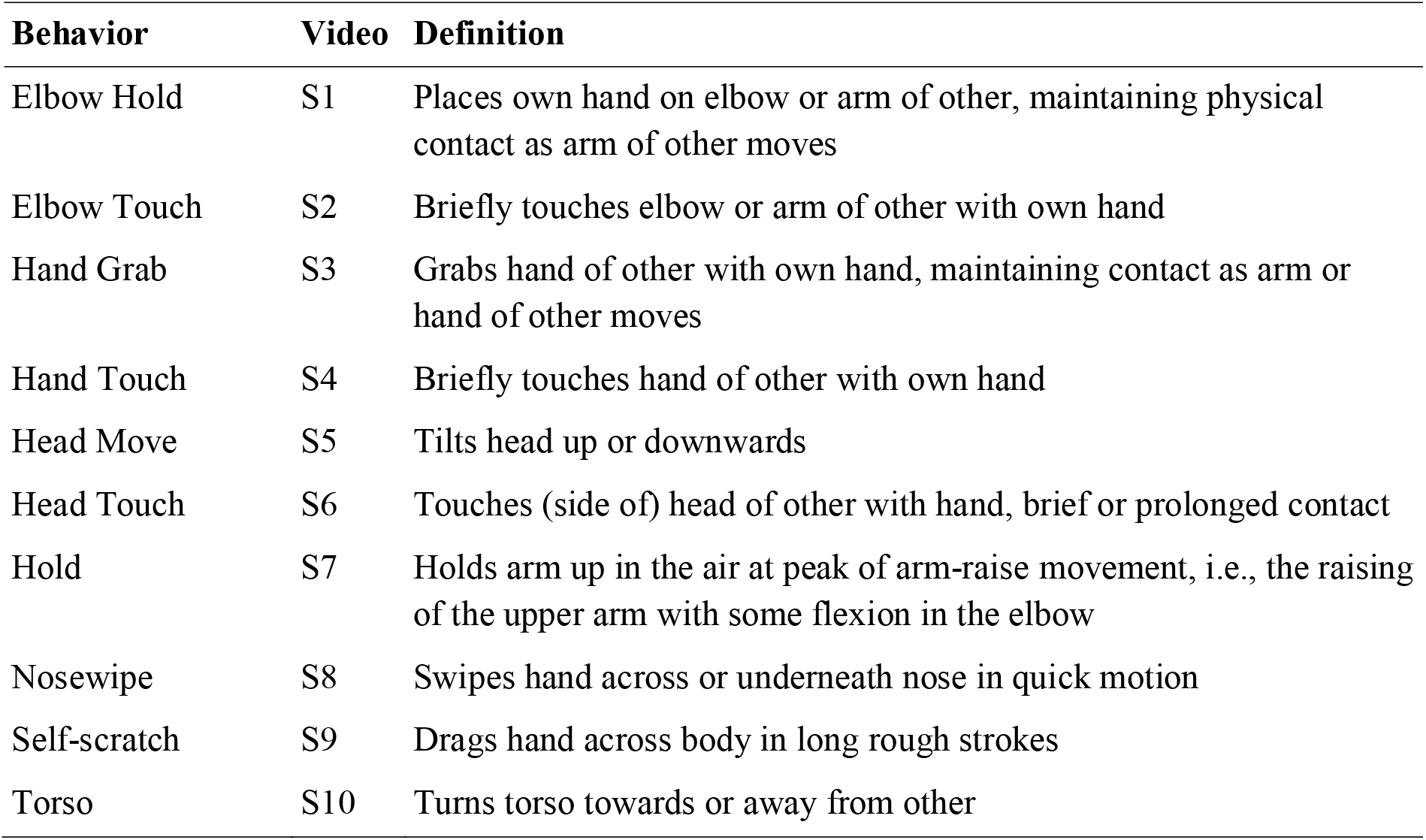
Ethogram of behavior, with reference to supplementary videos of behaviors.

**Table S3.**
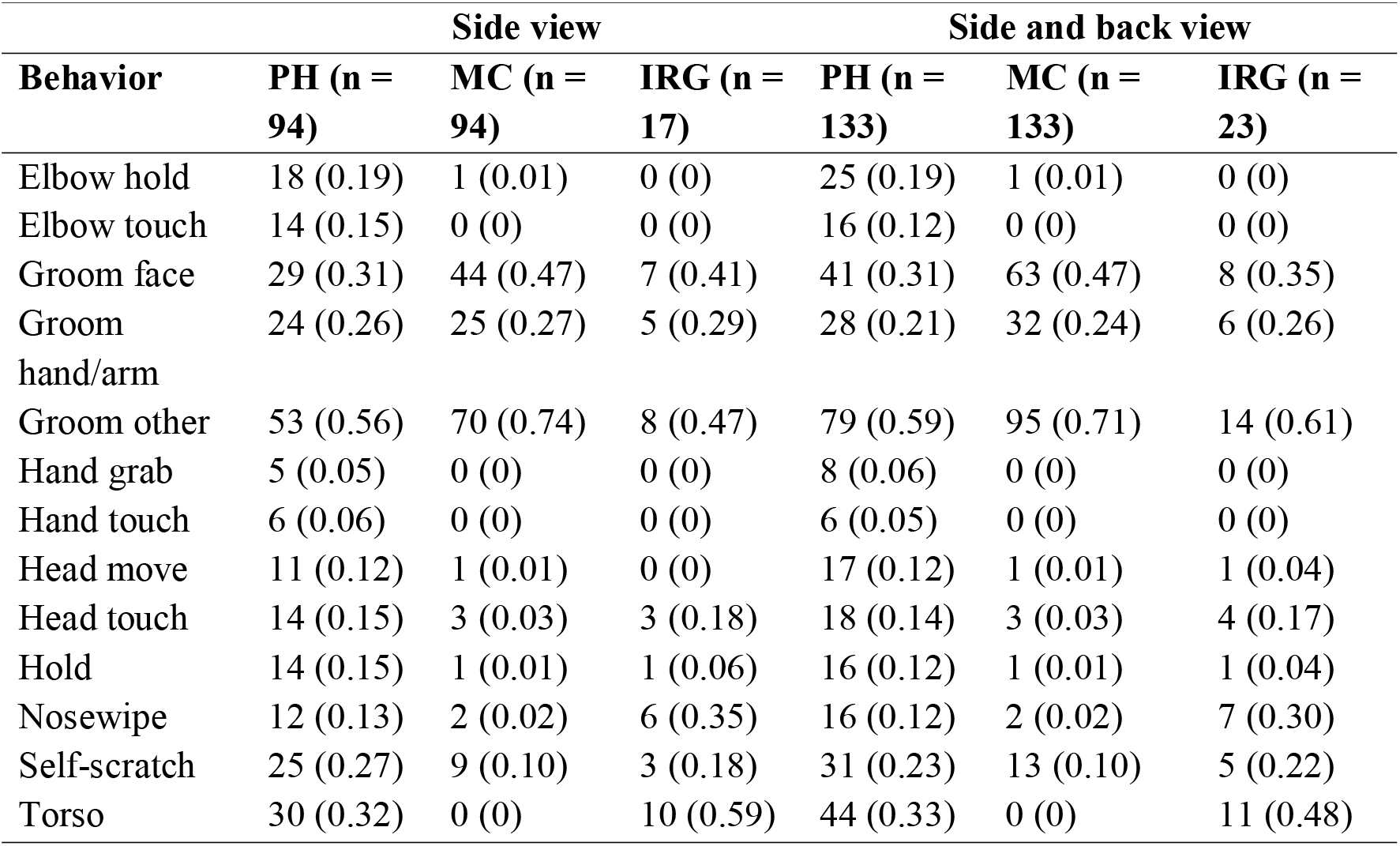
Occurrence of all ethogram behaviors in pre-handclasp (PH) and matched-control (MC) periods, as well as initiation of regular grooming bouts (IRG). Reported is the total number of instances and between brackets number of instances divided by number of bouts.

**Table S4.**
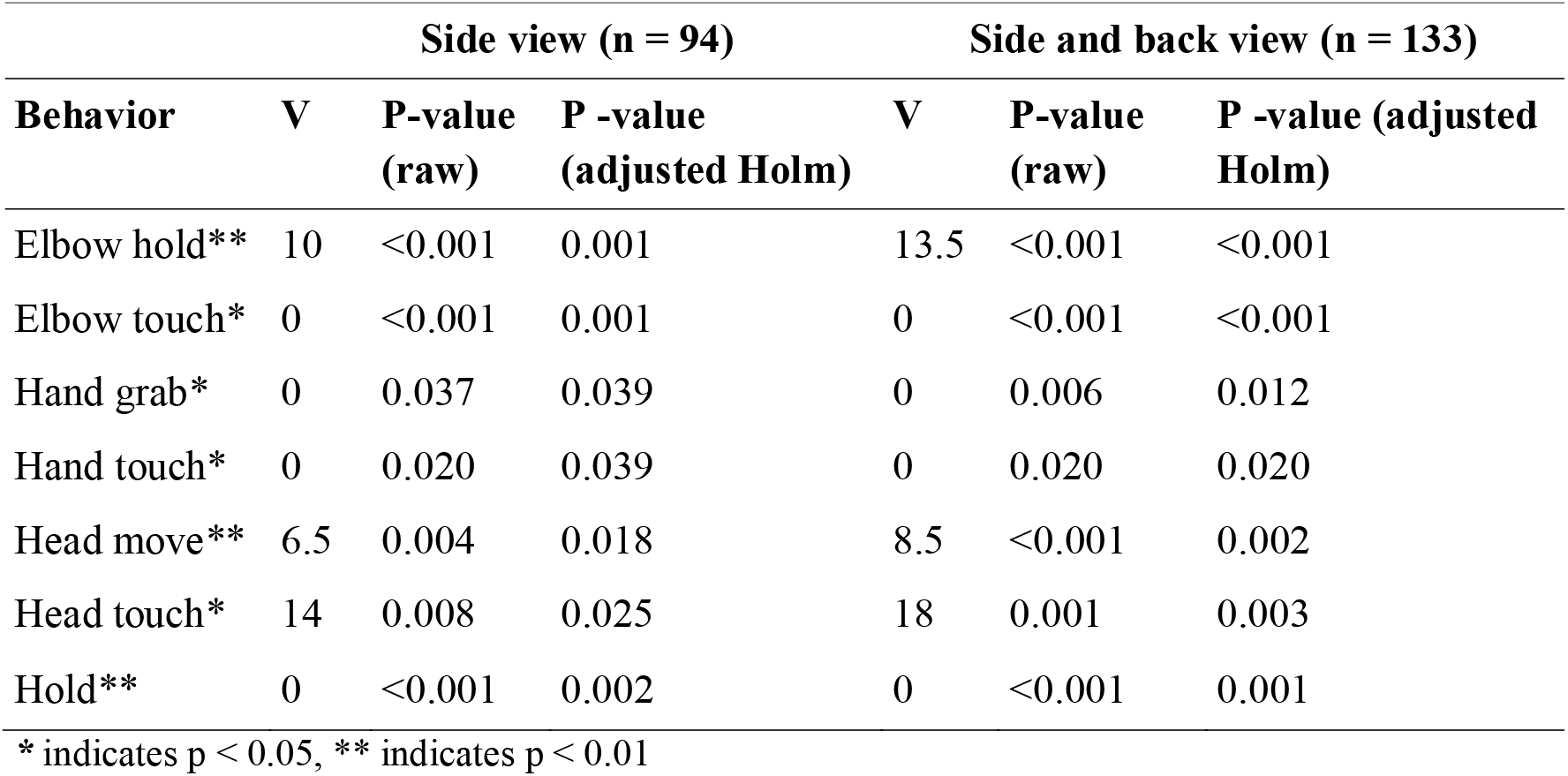
Results of the paired Wilcoxon signed rank test of behavior frequency between the PH and MC periods.

**Figure S14.**
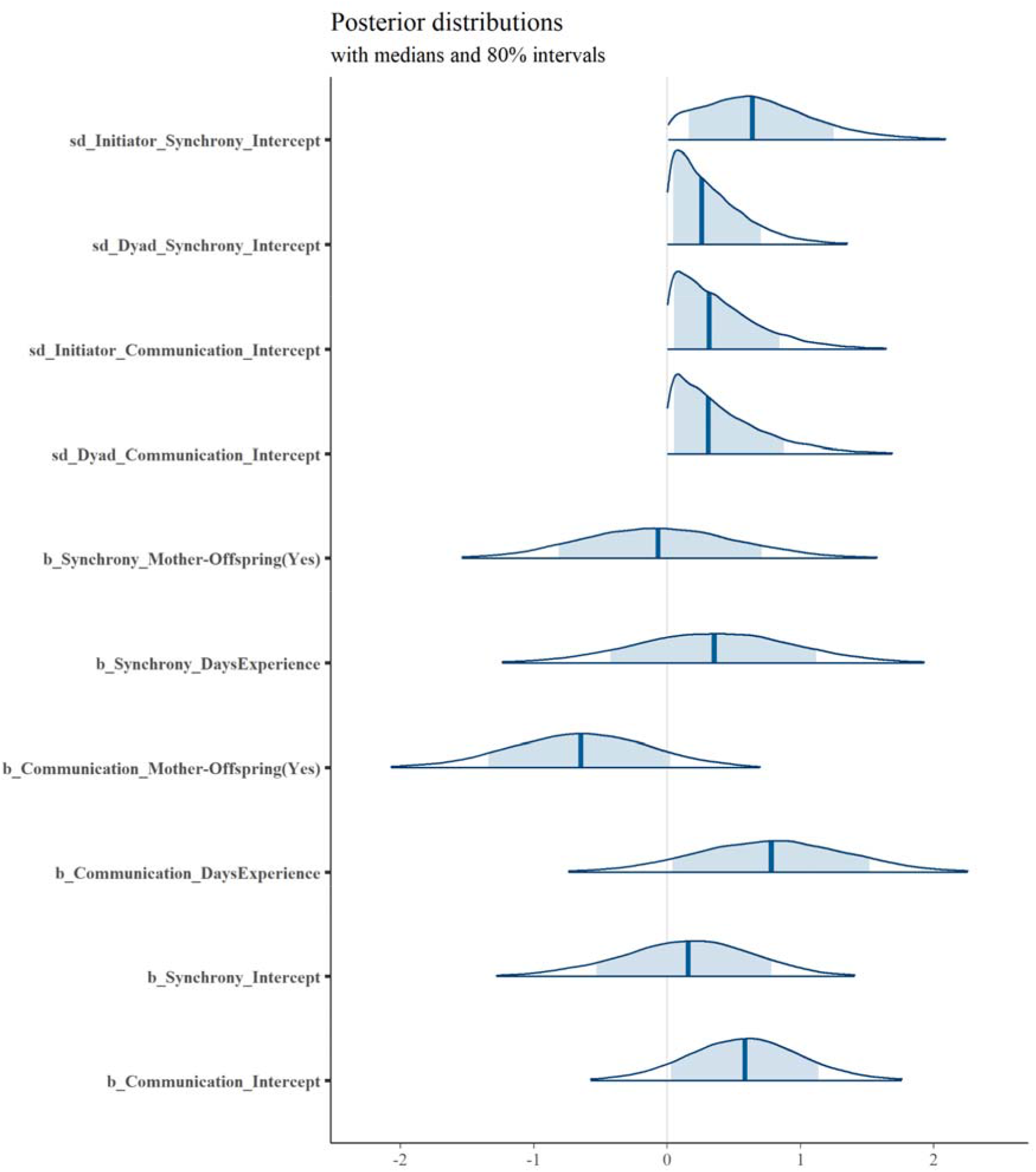
Posterior distributions of multinomial model exploring how type of GHC initiation (with shaping as reference category) was affected by dyadic days of experience with GHC and whether the dyad was mother-offspring or not. We included dyad and initiator ID as random effects, although for all but the intercept of initiator ID and synchrony the posterior distribution was near-identical to the prior. This indicates that the model could not reliably assess heterogeneity in GHC initiation type across dyad and individual identities with our data.

## Inter-rater reliability assessment

To establish inter-rater reliability, two additional coders (JB and AM) analyzed the PHs and MCs of 20% (N = 19) of the GHC bouts. Additionally, ZG coded this subset for a second time to compare to her first coding one year prior.

Due to the continuous nature of the behavioral scoring and the importance of observers not only agreeing on the presence of (often rare) behaviors but also the order in which they occurred, the usual Cohen’s Kappa approach was not feasible. We calculated inter-rater reliability as follows (see also attached data for each rater’s coding and an overview):

If raters both scored the same behavior in the same order at the same time (within 1 second) it was noted as an “agree”, if either of the raters scored a behavior the other one did not score it was a “disagree”. As an informed estimation for the total amount of behaviors per video we used the sum of the instances of agree and disagree. To correct for the probability of two raters agreeing on a behavior by chance, we multiplied the probability of scoring a particular behavior (1/11, the amount of different behaviors) by the probability of scoring it for a certain individual (0.5).

The final calculation of the reliability score combined this probability of agreement based on chance (POA) with the relative observer agreement (ROA, defined as agreed behaviors/total amount of behaviors), in the form of 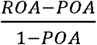. For instance, 50 agrees out of 70 total behaviors leads to a reliability score of 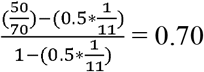. For this study, all dyads showed inter-rater reliability scores above 0.80 (ZG_2-ZG_1 = 0.89; JB-ZG_1 = 0.81; AM-ZG_1 = 0.82; JB-AM = 0.81). Furthermore, each rater also coded who they thought initiated the bout, and inter-rater reliability on initiator was calculated in a similar manner (with POA being 0.5) and was above 0.90 for all raters (JB-ZG_1 = 0.89; all other dyads = 1).

