## Supplemental Information for "Chimpanzees communicate to coordinate a cultural practice"

**Table S1.** Subject demography Group 2 at Chimfunshi Wildlife Orphanage on 01-07-2019.

|  | **Male** | **Female** | ***Total*** |
| --- | --- | --- | --- |
| **Adult (> 12 years)** | 8 | 24 | 32 |
| **Juvenile (3-11 years)** | 7 | 5 | 12 |
| **Infant (< 3 years)** | 3 | 5 | 8 |
| ***Total*** | 18 | 34 | 52 |

**Table S2.** Ethogram of behavior, with reference to supplementary videos of behaviors.

| **Behavior** | **Video** | **Definition** |
| --- | --- | --- |
| Elbow Hold | S1 | Places own hand on elbow or arm of other, maintaining physical contact as arm of other moves |
| Elbow Touch | S2 | Briefly touches elbow or arm of other with own hand |
| Hand Grab | S3 | Grabs hand of other with own hand, maintaining contact as arm or hand of other moves |
| Hand Touch | S4 | Briefly touches hand of other with own hand |
| Head Move | S5 | Tilts head up or downwards |
| Head Touch | S6 | Touches (side of) head of other with hand, brief or prolonged contact |
| Hold | S7 | Holds arm up in the air at peak of arm-raise movement, i.e., the raising of the upper arm with some flexion in the elbow |
| Nosewipe | S8 | Swipes hand across or underneath nose in quick motion |
| Self-scratch | S9 | Drags hand across body in long rough strokes |
| Torso | S10 | Turns torso towards or away from other |

**Table S3.** Occurrence of all ethogram behaviors in pre-handclasp (PH) and matched-control (MC) periods, as well as initiation of regular grooming bouts (IRG). Reported is the total number of instances and between brackets number of instances divided by number of bouts.

|  | **Side view** | | | **Side and back view** | | |
| --- | --- | --- | --- | --- | --- | --- |
| **Behavior** | **PH (n = 94)** | **MC (n = 94)** | **IRG (n = 17)** | **PH (n = 133)** | **MC (n = 133)** | **IRG (n = 23)** |
| Elbow hold | 18 (0.19) | 1 (0.01) | 0 (0) | 25 (0.19) | 1 (0.01) | 0 (0) |
| Elbow touch | 14 (0.15) | 0 (0) | 0 (0) | 16 (0.12) | 0 (0) | 0 (0) |
| Groom face | 29 (0.31) | 44 (0.47) | 7 (0.41) | 41 (0.31) | 63 (0.47) | 8 (0.35) |
| Groom hand/arm | 24 (0.26) | 25 (0.27) | 5 (0.29) | 28 (0.21) | 32 (0.24) | 6 (0.26) |
| Groom other | 53 (0.56) | 70 (0.74) | 8 (0.47) | 79 (0.59) | 95 (0.71) | 14 (0.61) |
| Hand grab | 5 (0.05) | 0 (0) | 0 (0) | 8 (0.06) | 0 (0) | 0 (0) |
| Hand touch | 6 (0.06) | 0 (0) | 0 (0) | 6 (0.05) | 0 (0) | 0 (0) |
| Head move | 11 (0.12) | 1 (0.01) | 0 (0) | 17 (0.12) | 1 (0.01) | 1 (0.04) |
| Head touch | 14 (0.15) | 3 (0.03) | 3 (0.18) | 18 (0.14) | 3 (0.03) | 4 (0.17) |
| Hold | 14 (0.15) | 1 (0.01) | 1 (0.06) | 16 (0.12) | 1 (0.01) | 1 (0.04) |
| Nosewipe | 12 (0.13) | 2 (0.02) | 6 (0.35) | 16 (0.12) | 2 (0.02) | 7 (0.30) |
| Self-scratch | 25 (0.27) | 9 (0.10) | 3 (0.18) | 31 (0.23) | 13 (0.10) | 5 (0.22) |
| Torso | 30 (0.32) | 0 (0) | 10 (0.59) | 44 (0.33) | 0 (0) | 11 (0.48) |

**Table S4.** Results of the paired Wilcoxon signed rank test of behavior frequency between the PH and MC periods.

|  | **Side view (n = 94)** | | | **Side and back view (n = 133)** | | |
| --- | --- | --- | --- | --- | --- | --- |
| **Behavior** | **V** | **P-value (raw)** | **P -value (adjusted Holm)** | **V** | **P-value (raw)** | **P -value (adjusted Holm)** |
| Elbow hold** | 10 | <0.001 | 0.001 | 13.5 | <0.001 | <0.001 |
| Elbow touch* | 0 | <0.001 | 0.001 | 0 | <0.001 | <0.001 |
| Hand grab* | 0 | 0.037 | 0.039 | 0 | 0.006 | 0.012 |
| Hand touch* | 0 | 0.020 | 0.039 | 0 | 0.020 | 0.020 |
| Head move** | 6.5 | 0.004 | 0.018 | 8.5 | <0.001 | 0.002 |
| Head touch* | 14 | 0.008 | 0.025 | 18 | 0.001 | 0.003 |
| Hold** | 0 | <0.001 | 0.002 | 0 | <0.001 | 0.001 |

******* indicates p < 0.05, ** indicates p < 0.01

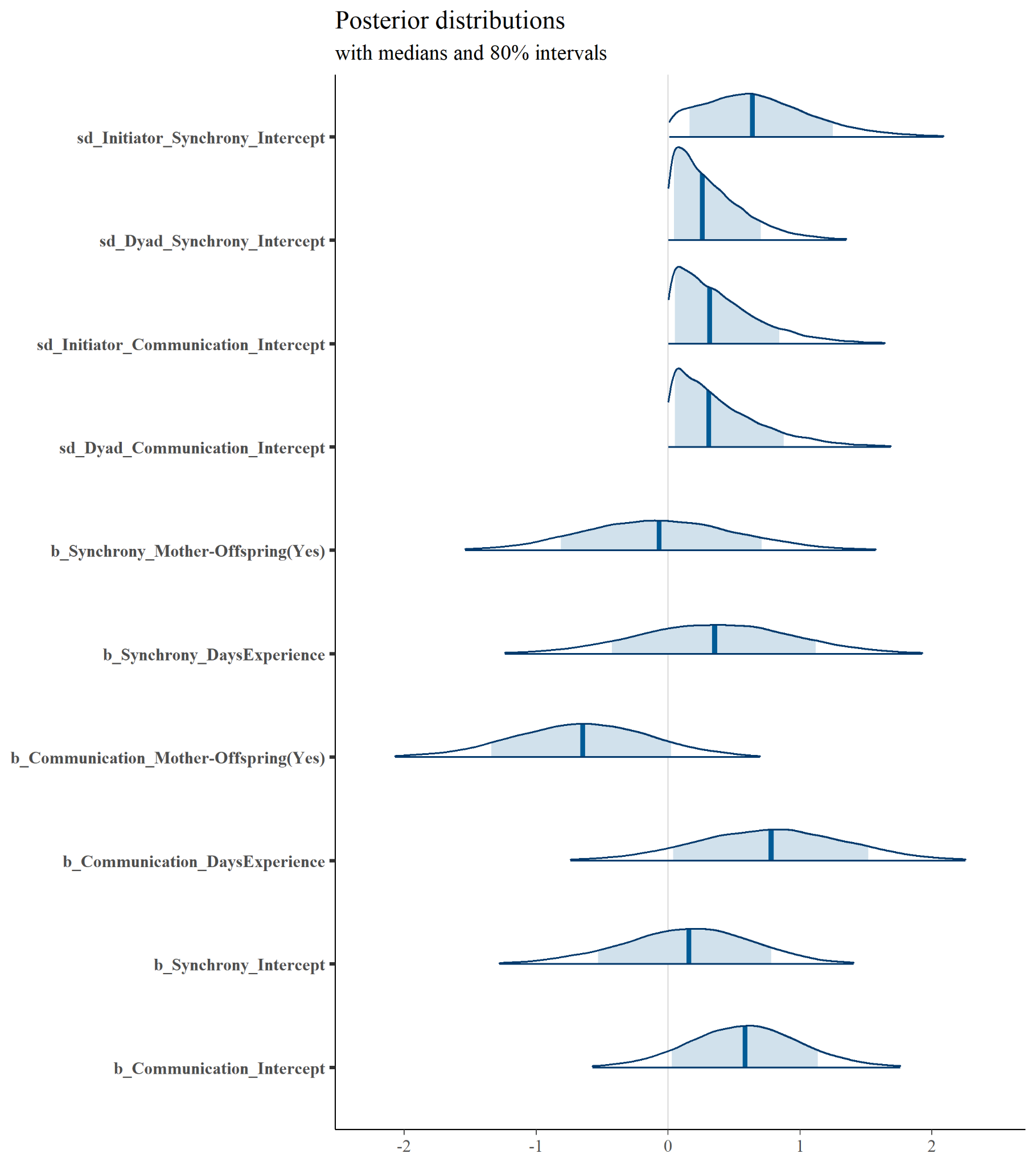

**Figure S14.** Posterior distributions of multinomial model exploring how type of GHC initiation (with shaping as reference category) was affected by dyadic days of experience with GHC and whether the dyad was mother-offspring or not. We included dyad and initiator ID as random effects, although for all but the intercept of initiator ID and synchrony the posterior distribution was near-identical to the prior. This indicates that the model could not reliably assess heterogeneity in GHC initiation type across dyad and individual identities with our data.
